# Genetic Deletion of Calcium-independent Phospholipase A_2_γ Protects Mice from Diabetic Nephropathy

**DOI:** 10.1101/2024.04.26.591364

**Authors:** Andrey V. Cybulsky, Joan Papillon, Julie Guillemette, José R. Navarro-Betancourt, Hanan Elimam, I. George Fantus

## Abstract

Calcium-independent phospholipase A_2_γ (iPLA_2_γ) is localized in glomerular epithelial cells (GECs)/podocytes at the mitochondria and endoplasmic reticulum, and can mediate release of arachidonic acid and prostanoids. Global knockout (KO) of iPLA_2_γ in mice did not cause albuminuria, but resulted in mitochondrial structural abnormalities and enhanced autophagy in podocytes. In acute glomerulonephritis, deletion of iPLA_2_γ exacerbated albuminuria and podocyte injury. This study addresses the role of iPLA_2_γ in diabetic nephropathy. Hyperglycemia was induced in male mice with streptozotocin (STZ). STZ induced progressive albuminuria in control mice (over 21 weeks), while albuminuria did not increase in iPLA_2_γ KO mice, remaining comparable to untreated groups. Despite similar exposure to STZ, the STZ-treated iPLA_2_γ KO mice developed a lower level of hyperglycemia compared to STZ-treated control. However, there was no significant correlation between the degree of hyperglycemia and albuminuria, and even iPLA_2_γ KO mice with greatest hyperglycemia did not develop significant albuminuria. Mortality at 21 weeks was greatest in diabetic control mice. Sclerotic glomeruli and enlarged glomerular capillary loops were increased significantly in diabetic control compared to diabetic iPLA_2_γ KO mice. Glomerular matrix was expanded in diabetic mice, with control exceeding iPLA_2_γ KO. Glomerular autophagy (increased LC3-II and decreased p62) was enhanced in diabetic iPLA_2_γ KO mice compared to control. Treatment of cultured GECs with H_2_O_2_ resulted in increased cell death in control GECs compared to iPLA_2_γ KO, and the increase was slightly greater in medium with high glucose compared to low glucose. H_2_O_2_-induced cell death was not affected by inhibition of prostanoid production with indomethacin. In conclusion, mice with global deletion of iPLA_2_γ are protected from developing chronic glomerular injury in diabetic nephropathy. This is associated with increased glomerular autophagy.

## Introduction

Among human glomerular diseases, diabetic nephropathy (DN) is a leading cause of chronic kidney disease, and has a major impact on health [1,2]. Current therapies of glomerulopathies and diabetic nephropathy are only partially effective, display significant toxicities and lack specificity [1]. Thus, it is important to understand the mechanisms of glomerular diseases to develop mechanism-based therapies.

Podocytes or glomerular visceral epithelial cells (GECs) are critical in the maintenance of glomerular permselectivity [3,4]. These cells display a complex morphology, characterized by cell bodies with projecting interdigitating foot processes that are bridged by filtration slit diaphragms. Their intricate shape is supported by the actin cytoskeleton. Podocytes are metabolically active cells with high energy demands; they produce slit-diaphragm proteins, adhesion molecules and glomerular basement membrane (GBM) components. Podocyte injury, which is characterized by albuminuria, is implicated in glomerular diseases, including diabetic nephropathy [3,4].

Calcium-independent phospholipase A_2_γ (iPLA_2_γ) catalyzes cleavage of fatty acids from the sn-1 or sn-2 position of phospholipids and from other substrates, such as cardiolipin [5]. We showed that iPLA_2_γ mRNA and protein are expressed in the glomerulus in vivo [6]. iPLA_2_γ is cytoprotective in complement-mediated GEC injury [6]. Furthermore, global knockout (KO) of iPLA_2_γ in mice results in striking mitochondrial ultrastructural abnormalities and enhances the number of autophagosomes in podocytes, leading to loss of podocytes in aging mice, but without detectable albuminuria [7]. However, there were no mitochondrial abnormalities or increased autophagosomes in mesangial and glomerular endothelial cells [7]. In experimental anti-GBM nephritis, and experimental focal segmental glomerulosclerosis (adriamycin nephrosis), deletion of iPLA_2_γ exacerbated albuminuria [7,8]. These studies thus demonstrated that iPLA_2_γ has a protective functional role in the normal glomerulus and in glomerulonephritis. In addition, our studies in cultured GECs verified that deletion of iPLA_2_γ is associated with mitochondrial dysfunction and enhanced autophagy [7].

We and others showed that iPLA_2_γ is localized subcellularly at the endoplasmic reticulum and mitochondria, and localization is dependent on the N-terminal region of iPLA_2_γ [5,9–11]. iPLA_2_γ can be active under resting and stimulated conditions [5]; complement-induced stimulation of iPLA_2_γ activity was dependent on phosphorylation at Ser-511 and/or Ser-515 via mitogen-activated protein kinase-interacting kinase 1 (MNK1) [10]. iPLA_2_γ can be activated by oxidative stress [12]. At the ER, iPLA_2_γ can modulate the unfolded protein response [13]. Phospholipases at the mitochondria are reported to play a key role in the regulation of mitochondrial function and signaling [5,11]. Global deletion of iPLA_2_γ in mice altered cardiolipin content and molecular species distribution that was accompanied by defects in mitochondrial function [14]. The role of iPLA_2_γ in mitochondrial bioenergetic function and its importance in cellular energy metabolism and homeostasis was demonstrated in several organs, including heart, skeletal muscle, adipose tissue, liver, and brain [14–18]. For example, iPLA_2_γ global KO mice display reduced growth rate, cold intolerance, and various bioenergetic dysfunctional phenotypes [15]. Global iPLA_2_γ deletion disrupted mitochondrial phospholipid homeostasis in the brains of aging mice, resulting in enlarged and degenerating mitochondria, enhanced autophagy and cognitive dysfunction [16]. Deletion of iPLA_2_γ in skeletal muscle resulted in muscle mitochondrial dysfunction and atrophy [19].

On the other hand, KO of iPLA_2_γ attenuated calcium-induced opening of the mitochondrial permeability transition pore and cytochrome c release [14]. Mice with myocardial KO of iPLA_2_γ showed less production of proinflammatory oxidized fatty acids, and developed substantially less cardiac necrosis compared with wild type littermates following ischemia-reperfusion injury [20]. In failing hearts, increased iPLA_2_γ activity channeled arachidonic acid into toxic hydroxyeicosatetraenoic acids (HETEs), promoting mitochondrial permeability transition pore opening, inducing myocardial necrosis/apoptosis and leading to further progression of heart failure [21]. Hepatic KO of iPLA_2_γ in mice subjected to high fat diet increased iPLA_2_γ-mediated hepatic 12-HETE production leading to mitochondrial dysfunction and hepatocyte death [22]. Deletion of iPLA_2_γ also reduced colonic tumorigenesis and venous thromboembolism [5]. Thus, iPLA_2_γ plays an important role in mitochondrial lipid metabolism and membrane structure, and perturbation of this role affects fatty acid β-oxidation, oxygen consumption, energy expenditure, and tissue homeostasis. However, the outcomes of iPLA_2_γ activity may be deleterious or protective depending on the context.

A small number of humans with mutations in the gene encoding iPLA_2_γ (PNPLA8) have been described. Phenotypes have included mitochondrial myopathy, lactic acidosis, microcephaly, seizures, neurodegeneration and ataxia; most cases have been severe, but two individuals survived into adulthood [23,24]. There is no information on renal phenotype.

The mechanisms and pathogenesis of diabetic nephropathy are complex and poorly understood [25–27]. Briefly, hyperglycemia and oxidative stress stimulate diacylglycerol, protein kinase C, the polyol pathway and advanced glycation end products, leading to protein glycosylation and stimulation of inflammatory mediators/cytokines and growth factors. This leads to renal hemodynamic changes and structural damage, in particular to podocytes, and podocyte injury is key in the pathogenesis of diabetic nephropathy [25,28–30]. A widely used experimental model of type 1 diabetes in rodents is induced by streptozotocin (STZ), a chemical toxin for pancreatic β-cells. Renal injury resembles human diabetic nephropathy, and it includes albuminuria, podocyte loss, mesangial and GBM expansion, decline in kidney function. Eventually, this progresses to glomerular and tubulointestitial sclerosis, although a limitation of the model is that injury is relatively mild [27,28,31]. In the present study, we address the role of iPLA_2_γ in diabetic nephropathy – a clinically-important cause of glomerular injury, whose pathogenesis is distinct from the primary/acute glomerulopathies. In contrast to acute glomerulonephritis, mice with global deletion of iPLA_2_γ are protected from developing chronic glomerular injury in diabetic nephropathy.

## Materials and methods

### Antibodies and chemicals

Goat anti-synaptopodin antibody (sc-21537) was purchased from Santa Cruz Biotechnology (Santa Cruz, CA). Rabbit anti-Wilms tumor-1 (WT1) antibody (CAN-R9(IHC)-56-2; ab89901) was from Abcam Inc. (Toronto, ON). Rat anti-collagen α5 (IV) antibody clone H53 (7078) was purchased from Chondrex Inc. (Woodinville, WA). Rabbit antibodies to LC3B (2775) and SQSTM1/p62 (5114) were purchased from Cell Signaling Technology (Danvers, MA). Mouse anti-LC3B clone 5F10 (ALX-830-080-C100) was from Enzo Life Sciences (Ann Arbor, MI). Rabbit anti-ubiquitin (U5379) and rabbit anti-actin (A2066) antibodies were from MilliporeSigma (Mississauga, ON). Rat anti-mouse F4/80 antibody MCA497GA was from Bio-Rad (Mississauga, ON). Goat anti-podocalyxin antibody (AF1556) was purchased from R & D Systems (Minneapolis, MN). Rabbit anti-nephrin antiserum was a gift from Dr. Tomoko Takano (McGill University) [32]. Non-immune IgG and secondary antibodies were from Jackson ImmunoResearch Laboratories (West Grove, PA) or Thermo-Fisher Scientific. Streptozotocin (STZ) and rhodamine-phalloidin were from MilliporeSigma. 5(6)-Carboxy-2′,7′-dichlorofluorescein diacetate N-succinimidyl ester (DCF) was from Santa Cruz Biotechnology.

### Studies in mice (in vivo)

iPLA_2_γ KO mice in a C57BL/6 background were kindly provided by Dr. Richard Gross (Washington University, St. Louis, MO, USA). Mice were produced, bred and genotyped, as described previously [7,15]. The animals were housed in a pathogen free facility, under standard conditions including cages and bedding, with 12 h on-off light cycles, and were fed ad libitum. Adult (6 month old) male control and iPLA_2_γ KO littermates were untreated or received STZ 50 mg/kg (in 50 mM sodium citrate buffer, pH 4.5) intraperitoneally daily for 5 days [31]. Mice were given 10% sucrose water to drink for 6 days after STZ treatment to prevent fatal hypoglycemia. The animals were randomly allocated to experimental groups. Blood glucose was measured after 1 week using glucose test strips. If hyperglycemia did not develop, the protocol, was repeated. This low-dose STZ protocol avoids direct STZ renal toxicity. Mice were followed for ∼6 months and were euthanized (isofluorane followed by cervical dislocation) in the animal facility to collect the kidneys and isolate glomeruli by sequential sieving [7]. The animal protocol was approved by the McGill University Animal Care Committee, and our studies comply with the guidelines established by the Canadian Council on Animal Care. The study complied with ARRIVE guidelines.

Urine collections were performed in the morning in the animal facility at monthly intervals until the mice were euthanized. Urine albumin was quantified with an enzyme-linked immunosorbent assay (Mouse Albumin ELISA Quantification Kit, Bethyl Laboratories, Montgomery, TX). Albumin results were normalized to urine creatinine, which was measured using a picric acid-based reaction (Creatinine Colorimetric Assay Kit, Cayman Chemical Co; Ann Arbor, MI) [7,32].

### Studies in cell culture

Primary GECs were derived from iPLA_2_γ KO mice and wild type control mice. The detailed method and characterization of the cells was published previously [13]. Control and iPLA_2_γ KO cell lines were cultured on plastic substratum in K1 medium (DMEM, Ham F-12, with 5% NuSerum and hormone mixture). For experiments, GECs were allowed to proliferate for one day at 33°C and were then switched to 37°C to differentiate for 24 h. The lactate dehydrogenase (LDH) release assay to assess cell death was described previously [6,7].

### Microscopy

For light microscopy, portions of kidneys were fixed in 4% paraformaldehyde and stained with periodic acid-Schiff by conventional techniques at the McGill University Health Centre Histology Platform. To minimize observer bias, quantitative morphometry was used to characterize histological changes. Slides were digitized at 40x resolution in an Aperio AT Turbo scanner (Leica Biosystems, Buffalo Grove, IL). Images were processed using Aperio ImageScope 12.4 (Leica Biosystems). Glomeruli in experimental groups were selected randomly and analyzed with the Positive Pixel Count v9 algorithm, as reported previously [32]. Positive pixels were identified by a hue value of 0.854 (pink) and a hue width of 0.035. Glomerular matrix expansion was expressed as the ratio of positive over total pixels. To quantify podocyte number (WT1 counts) and kidney macrophages (F4/80 staining), kidney sections were deparaffinized and rehydrated. Antigen retrieval was done using citrate buffer, pH 6. Automated immunohistochemistry staining was performed by the McGill University Health Centre Histology Platform, using the Discovery Ultra Instrument (Roche Diagnostics). For WT1, the WT1-positive nuclei were then quantified by visual counting.

For immunofluorescence (IF) microscopy, kidney poles were snap-frozen in isopentane (−80°C). Cryostat sections (4 μm thickness) were cut and then fixed in 4% paraformaldehyde (22°C), ice-cold methanol or ice-cold acetone, and blocked with 5% normal rabbit or goat serum or 3-5% BSA. Incubations with primary antibodies were performed overnight at 4°C, and incubations with secondary antibodies were 1 h at 22°C. In control incubations (performed in parallel), primary antibody was replaced with nonimmune IgG. In some experiments, cell nuclei were stained with Hoechst H33342 [32,33]. Glomeruli in experimental groups were selected randomly and images were acquired using a Zeiss Axio Observer Z1 LSM780 laser scanning confocal microscope with ZEN2010 software (McGill University Health Centre Research Institute Imaging Platform). To compare fluorescence intensities, all images were taken at the same exposure time. Fluorescence intensity was quantified using the histogram function of ImageJ software (National Institutes of Health, Bethesda, MD), and results are expressed in arbitrary units [32,33]. The glomerular fluorescence intensity was normalized to the total fluorescence in each image. To measure the colocalization of two proteins in mouse kidney sections, glomeruli were circled, and threshold intensity of each channel was measured, as previously. Colocalization of the thresholded images in kidney sections was determined using the JACoP plugin in ImageJ [33].

### Immunoblotting

The immunoblotting protocol was described previously [7,32]. Chemiluminescence was detected in a ChemiDoc Touch Imaging System (Bio-Rad; Mississauga, ON). Signal saturation was monitored with Image Lab (Bio-Rad) and only signal intensities within a linear range were analyzed. Densitometry of bands was quantified using ImageJ, and values were normalized to the expression of β-actin.

### Statistical analysis

Values are expressed as mean ± standard error. Data were processed in Prism (GraphPad Software, La Jolla, CA). In experiments with three or more groups, or groups and multiple time-points, one-way or two-way analysis of variance (ANOVA) was used to determine significant differences among groups; where relevant, additional comparisons were calculated and values were adjusted according to the Holm-Sidak method. Significant differences between two groups are displayed with lines between columns, with asterisks denoting P values. In the absence of such lines, differences were not statistically significant. Other statistical tests are presented in the figure legends.

## Results

### KO of iPLA_2_γ attenuates development of albuminuria in STZ-diabetic nephropathy

Mice with global deletion of iPLA_2_γ (mean age 6.5 months) were employed to address the functional role of iPLA_2_γ in STZ-induced diabetic nephropathy. Within one month after treatment of mice with STZ, increases in blood glucose were evident in both control and iPLA_2_γ KO mice, indicating development of diabetes (Supplementary Fig. S1A). STZ induced significant increases in glucose in both control and iPLA_2_γ KO mice; however, mean glucose levels over the study period were greater in control mice. Adult iPLA_2_γ KO mice are known to be smaller than wild type mice [15,16,22]. In keeping with this, untreated and STZ-treated iPLA_2_γ KO mice showed lower body weights compared to untreated and STZ-treated controls. Furthermore, STZ-treated control mice showed lower body weight compared to untreated control, whereas the weight of STZ-treated iPLA_2_γ KO mice did not decrease further below the already smaller untreated iPLA_2_γ animals (Supplementary Fig. S1C).

Induction of diabetes induced progressive albuminuria in control mice, and the albuminuria in diabetic control mice was significantly greater compared to diabetic iPLA_2_γ KO mice (the week 2-21 time points were considered together in the analysis). Diabetic iPLA_2_γ KO mice showed slightly although not significantly greater albuminuria compared to untreated iPLA_2_γ KO mice, and the levels of albuminuria in these KO mice were comparable to untreated control (Fig. 1A).

**Figure 1.**
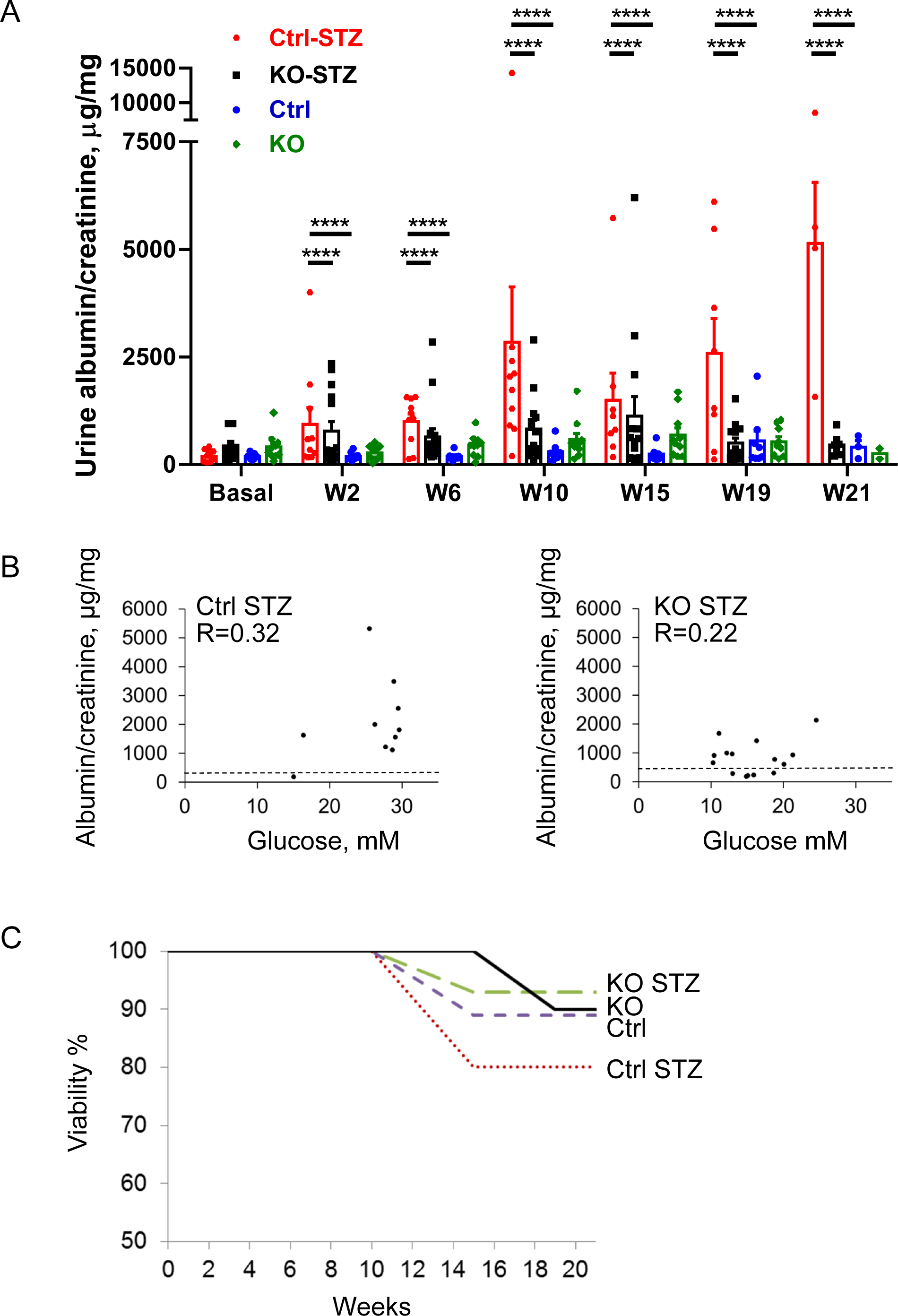
KO of iPLA_2_γ attenuates development of albuminuria in STZ-induced diabetic nephropathy. A) Treatment of mice with STZ induces progressive albuminuria (monitored as urine albumin/creatinine) in control mice. N=9 mice in control (Ctrl) untreated (Untr), 10 in KO Untr, 10 in Ctrl STZ and 15 in KO STZ groups. ****P<0.0001 Ctrl STZ vs KO STZ, P<0.0001 Ctrl STZ vs Ctrl untreated. KO Untr vs KO STZ is not significant (the week 2-21 time points were considered together in the analysis; two-way ANOVA). W, week. B) Correlation of albuminuria with blood glucose. The mean week 2-21 albumin/creatinine value of each diabetic mouse are plotted against mean blood glucose value of the same mouse. The correlations in each group are relatively weak. The dashed lines indicate the mean urine albumin/creatinine values of the respective untreated groups. C) Viability of mice. Greatest mortality is evident in the Ctrl STZ mice.

Since blood glucose levels were greater in diabetic control mice compared to iPLA_2_γ KO, we removed 7 mice from analysis of the iPLA_2_γ KO group. This resulted in an increase in the mean blood glucose in the remaining subgroup of diabetic iPLA_2_γ KO mice (Supplementary Fig. S1B); however, the mean urine albumin/creatinine values at all time points did not change significantly and remained substantially lower compared to diabetic controls (Supplementary Fig. S2). Indeed, there was a weak correlation between the mean albumin/creatinine value of each diabetic mouse and mean blood glucose value of the same mouse in both the control and iPLA_2_γ KO groups (Fig. 1). Finally, the greatest mortality was evident in the diabetic control mice (Fig. 1).

### Diabetic control mice show alterations in glomerular morphology

Mouse kidneys were isolated after 21 weeks of diabetes. Both diabetic control and iPLA_2_γ KO mice showed increases in glomerular matrix, compared to the respective untreated groups. Furthermore, the increase was significantly greater in control mice (Fig. 2). Glomerular cross-sectional area was not substantially different among groups of mice, although statistically, it was reduced slightly in diabetic iPLA_2_γ KO mice compared to untreated iPLA_2_γ KO (Fig. 2).

**Figure 2.**
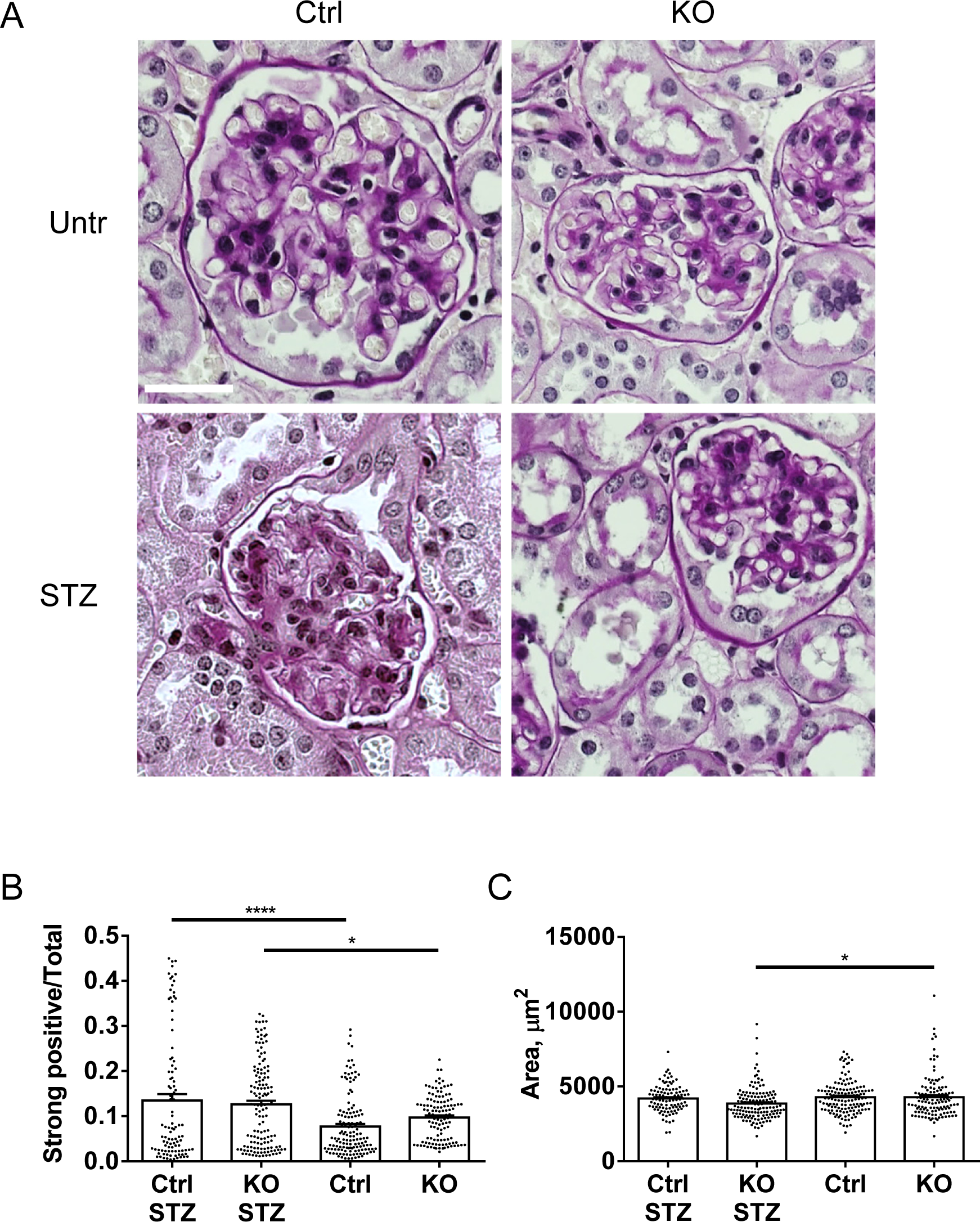
STZ increases glomerular matrix expansion. Kidney sections were stained with periodic acid-Schiff and glomerular matrix expansion was evaluated with a pixel-counting algorithm. A) Representative photomicrographs. B) Quantification of extracellular matrix. Both STZ-treated control and iPLA_2_γ KO mice show significant increases in glomerular matrix, compared to untreated groups, and the increase is greater in control (P<0.02). C) Glomerular cross-sectional area is reduced slightly in iPLA_2_γ KO STZ compared to KO untreated mice and tends to be slightly lower in control STZ compared to control untreated. 20 glomeruli in 5-6 mice per group were analyzed. *P<0.05, ****P<0.0001 (ANOVA). Bar = 25 µm.

Kidney sections were immunostained with antibodies to the podocyte protein synaptopodin and collagen IV-α5. Importantly, diabetic control mice showed an increase in sclerotic glomeruli, and although there was no change in glomerular area, the diabetic control glomeruli showed large capillary loops, compared to the other groups (Fig. 3). The latter is compatible with reduced glomerular contractility and glomerular hypertension/hyperfiltration.

**Figure 3.**
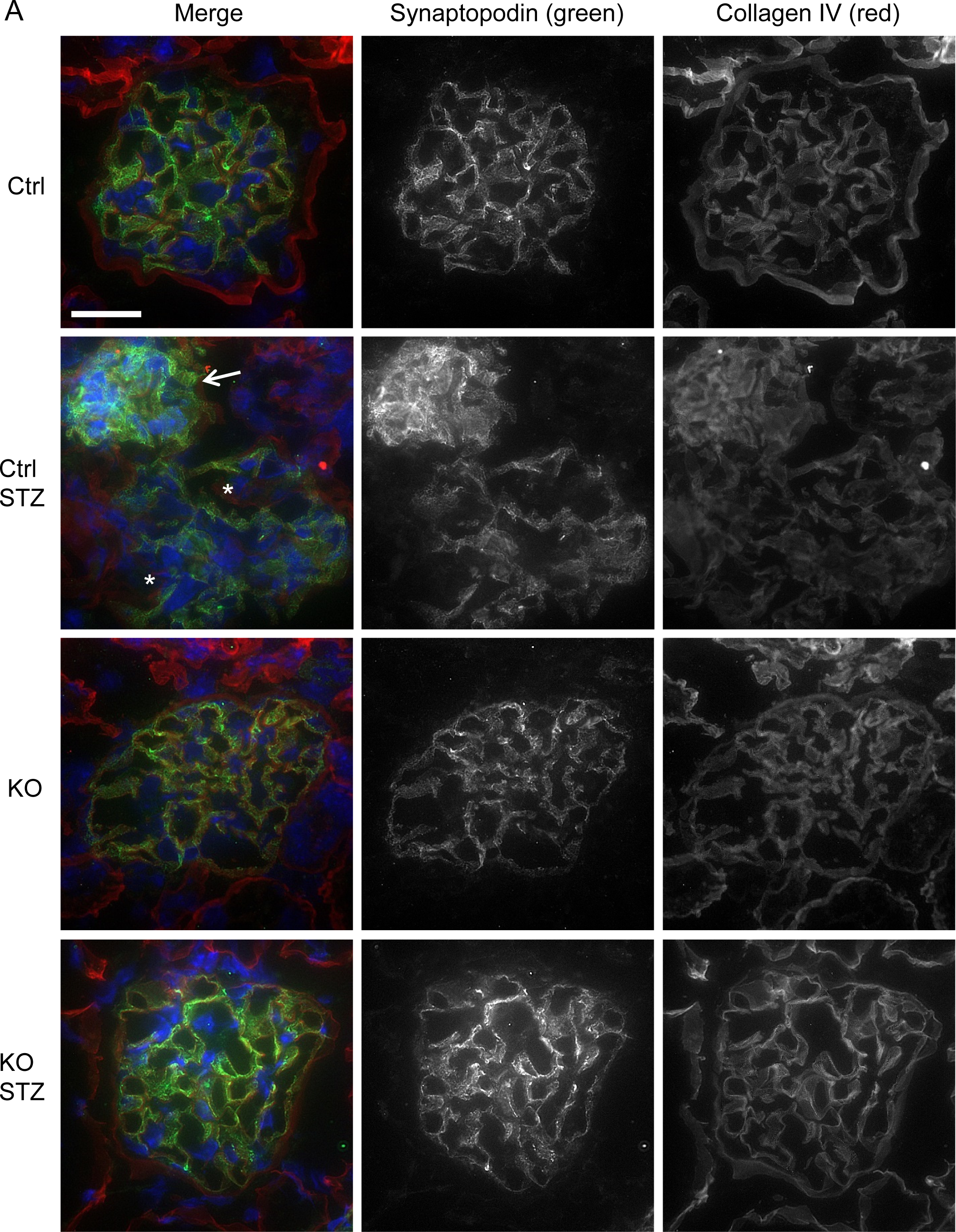
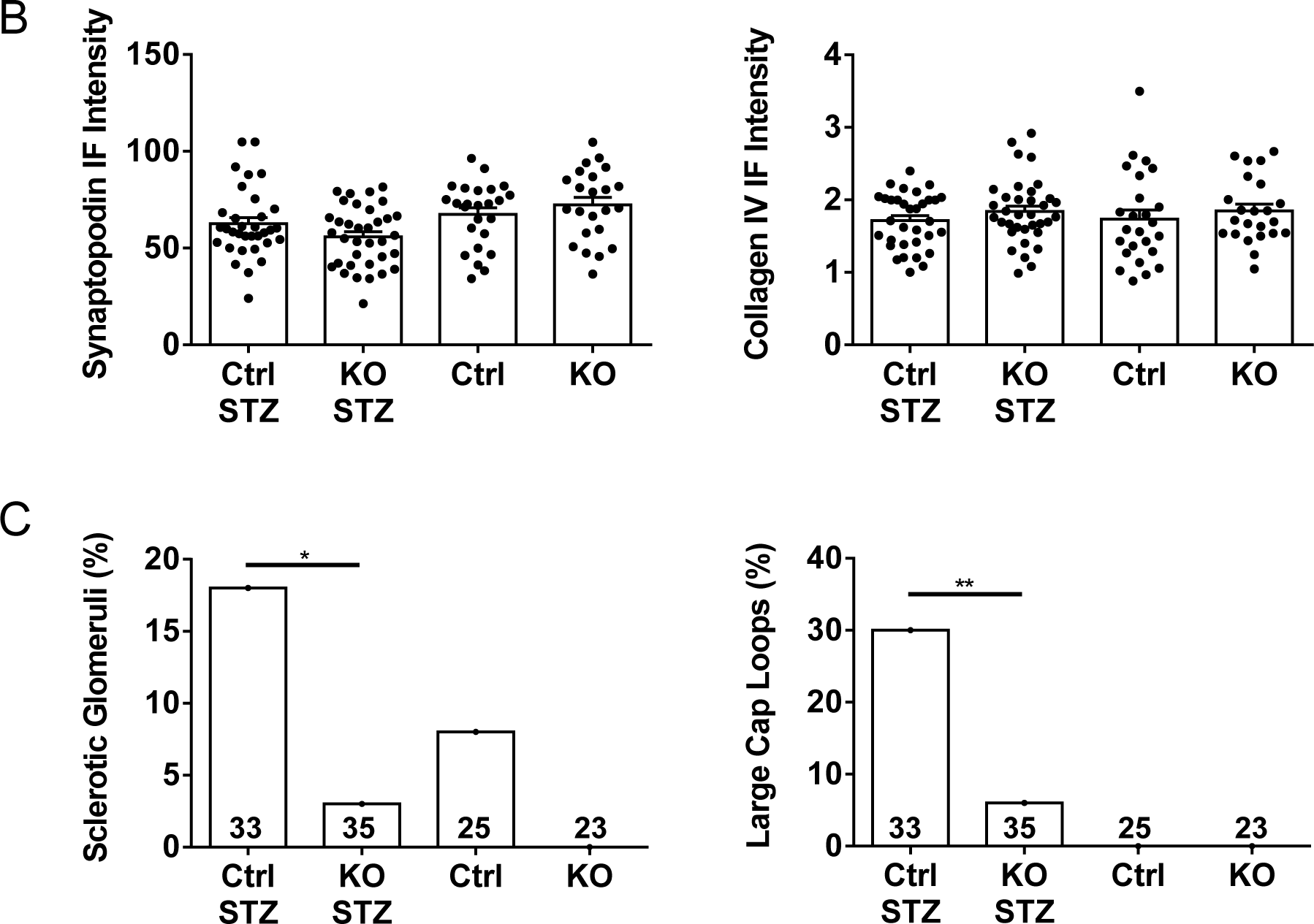
Control (Ctrl) STZ mice show changes in glomerular morphology. A) Kidney sections were stained with antibodies to synaptopodin and collagen IV-α5 (representative immunofluorescence micrographs). B) Quantification of immunofluorescence intensity. There are no significant differences in synaptopodin or collagen immunofluorescence intensities among groups (ANOVA). However, Ctrl STZ mice show an increase in sclerotic glomeruli (A; arrow) and glomeruli with large capillary loops (*) compared to the other groups (quantification in panel C). 5-7 glomeruli/mouse in 4-6 mice per group were analyzed. *P<0.05, **P<0.01 (Chi-squared test). Bar = 25 µm.

The presence of albuminuria in diabetic control mice reflects podocyte injury. However, we did not observe a loss of podocytes, i.e. there were no significant differences in WT1 counts (podocyte numbers) or WT1/glomerular area counts among groups (Fig. 4). In addition, there were no significant differences in the podocyte protein synaptopodin or glomerular collagen IV-α5 immunofluorescence staining intensities among groups (Fig. 3). Colocalization of synaptopodin and collagen IV-α5 was not substantially different among groups. The Pearson correlation coefficients were 0.67±0.05 (untreated control), 0.72±0.05 (untreated iPLA_2_γ KO), 0.69±0.08 (diabetic control) and 0.66±0.04 (diabetic iPLA_2_γ KO). These values imply that most of the collagen IV-α5 was localized in or near podocytes. Finally, glomeruli were isolated from mouse kidneys and lysates were subjected to immunoblotting to examine expression of nephrin and podocalyxin. There were no significant differences in the expression of these proteins among groups (Supplementary Fig. S3).

**Figure 4.**
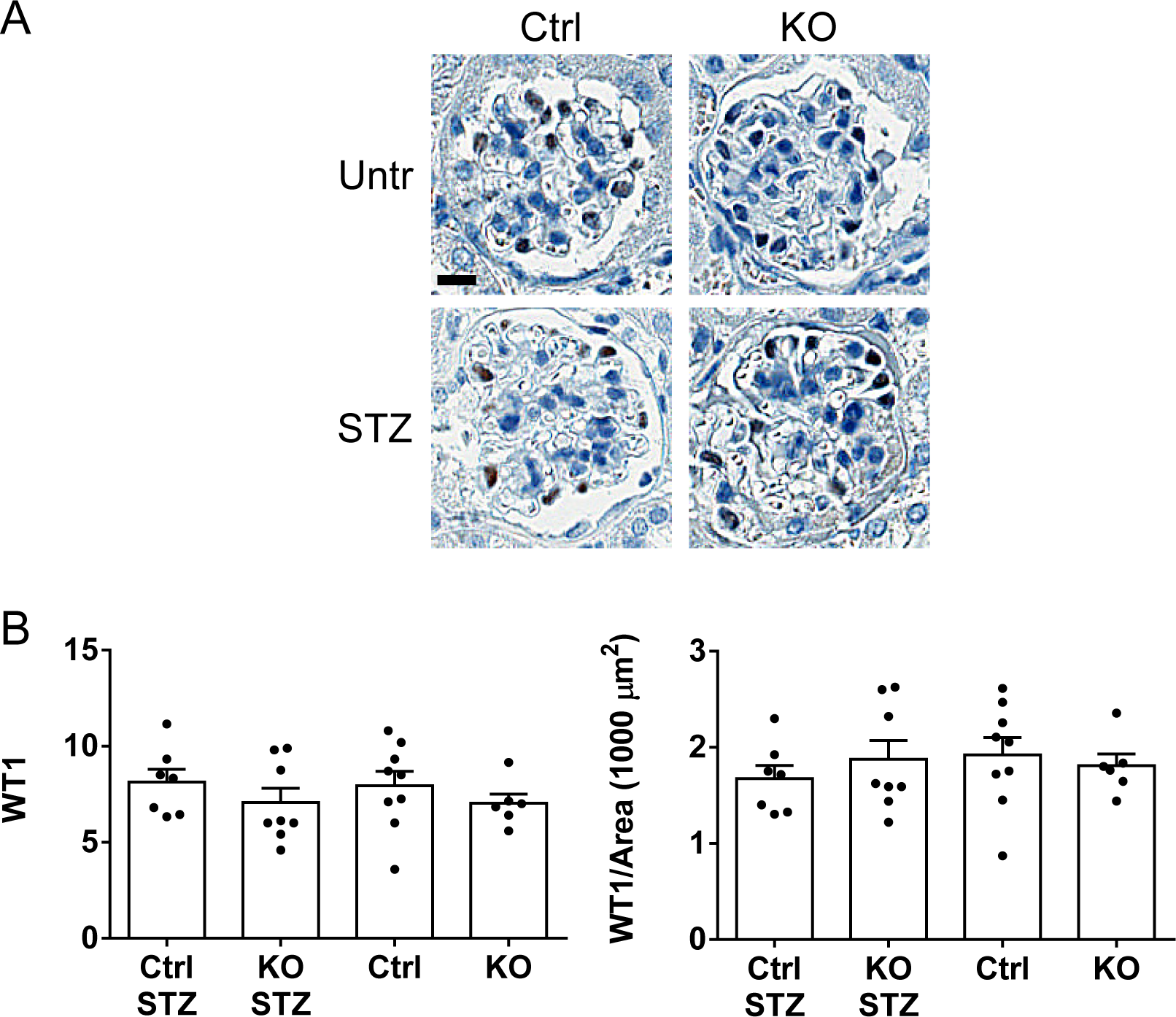
Podocyte numbers in control and iPLA_2_γ KO mice (WT1-positive cells). A) Kidney sections were stained with anti-WT1 antibody (representative photomicrographs). B) WT1 counts. There are no significant differences in WT1 counts or WT1/glomerular area counts among groups (ANOVA). 10 glomeruli/mouse in 5-7 mice per group were analyzed; each point is the mean value of a single mouse. Bar = 25 µm.

The complex structure of podocytes, in particular the foot processes is dependent on the actin cytoskeleton [3,4]. To examine the effect of iPLA_2_γ and diabetes on actin cytoskeleton organization in podocytes, kidney sections were stained with phalloidin (which labels F-actin) and synaptopodin [34]. Confocal fluorescence microscopy demonstrated substantial colocalization of F-actin and synaptopodin (Pearson colocalization coefficient >0.75). In keeping with our findings on WT1 counts and expression of podocyte proteins, glomerular F-actin content (phalloidin fluorescence intensity) and F-actin colocalization with synaptopodin did not show significant differences among the groups of mice, implying that deletion of iPLA_2_γ and diabetes did not substantially affect cytoskeleton organization and foot processes (Supplementary Fig. S4).

### Inflammatory cells in diabetic kidneys

Diabetic nephropathy may represent a proinflammatory condition associated with infiltration of inflammatory cells [25,35]. To examine for macrophages, kidney sections were stained with the F4/80 macrophage-specific antibody. There was only minimal F4/80 staining in the 4 groups of mice, suggesting that macrophages were largely absent (Supplementary Fig. S5). Periodic acid-Schiff-stained slides were also examined for the presence of neutrophils. Occasional neutrophils were visualized in the kidney sections, but the numbers were very small and did not allow for accurate quantification (Supplementary Fig. S6).

### Autophagy is enhanced in iPLA_2_γ KO mice

Protein misfolding and impaired clearance of misfolded proteins in glomerular cells, particularly podocytes may contribute to podocyte injury in glomerulopathies [36]. In this set of experiments, we examined the effects of iPLA_2_γ and diabetes on pathways of protein degradation. Among these, autophagy is a prominent pathway that regulates protein homeostasis in podocytes [36]. Lysates of glomeruli isolated after 21 weeks of diabetes were immunoblotted with anti-LC3 or anti-p62 antibodies [7,8,32]. LC3-II, a marker of autophagy, was increased significantly in untreated and diabetic iPLA_2_γ KO mice compared to untreated and diabetic control (Fig. 5). p62 (an autophagy substrate) was decreased in diabetic iPLA_2_γ KO mice compared to the other groups (Fig. 5), implying enhanced degradation of p62 by autophagy in the diabetic iPLA_2_γ KO mice. Thus, autophagy was enhanced in diabetic iPLA_2_γ KO mice, while untreated iPLA_2_γ KO may have higher basal autophagy. In the diabetic iPLA_2_γ KO mice, autophagy correlated with lower albuminuria, and could thus represent a protective pathway.

**Figure 5.**
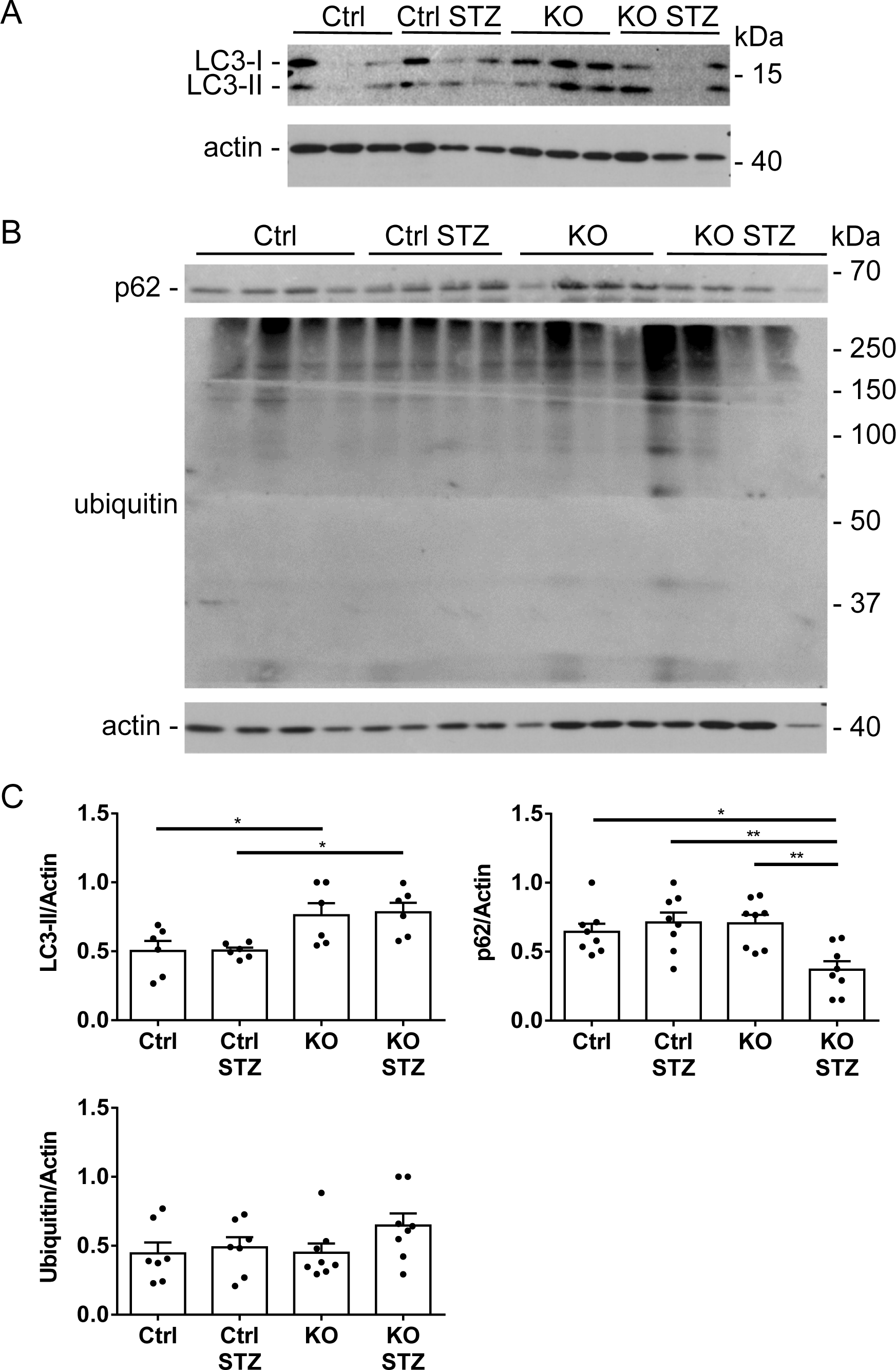
iPLA_2_γ KO mice show increased glomerular autophagy. A and B) Glomerular lysates were immunoblotted with antibodies to LC3, p62 and ubiquitin (representative immunoblots). C) Signals were quantified by densitometry. LC3-II is increased significantly in untreated and STZ-treated iPLA_2_γ KO mice compared to untreated and STZ-treated control. p62 is decreased significantly in STZ-treated iPLA_2_γ KO mice. There are no significant differences in protein ubiquitination among groups. There are 6-8 mice per group. *P<0.05, **P<0.01 (ANOVA).

To address protein degradation via the ubiquitin-proteasome system [8,36], lysates were immunoblotted with anti-ubiquitin antibody. In contrast to LC3-II, there were no significant differences in protein ubiquitination among groups (Fig. 5), suggesting there were no substantial differences in the function of the ubiquitin-proteasome system.

### Control GECs are more susceptible to injury compared to iPLA_2_γ KO GECs

As discussed above, diabetic control mice showed greater albuminuria compared to iPLA_2_γ KO mice (Fig. 1). Therefore, we examined if cultured GECs derived from control mice are more susceptible to injury compared to GECs from iPLA_2_γ KO mice. GECs were untreated, or treated with potentially cytotoxic compounds, including tunicamycin, adriamycin or H_2_O_2_. To reflect the diabetic environment in vivo, experiments were carried out in medium with a high glucose concentration (36 mM); low glucose medium was used as control (7.8 mM). By analogy to an approach employed in hepatocytes [22], injury was assessed by monitoring release of LDH from cells into cell supernatants. Control GECs were more susceptible to adriamycin- and H_2_O_2_-induced cytolysis compared to iPLA_2_γ KO GECs (Fig. 6A), thus recapitulating injury in vivo. Furthermore, in control GECs, H_2_O_2_-induced cytolysis was greater in high glucose medium compared to low glucose. Adriamycin-induced cytolysis in control GECs was slightly greater in high glucose medium compared to low glucose, but the difference did not reach statistical significance. Similarly to H_2_O_2_, adriamycin induces cytotoxicity via production of reactive oxygen species, but adriamycin can also activate other cytotoxic pathways that are less dependent on a high glucose milieu. Tunicamycin was not cytotoxic at the doses employed.

**Figure 6.**
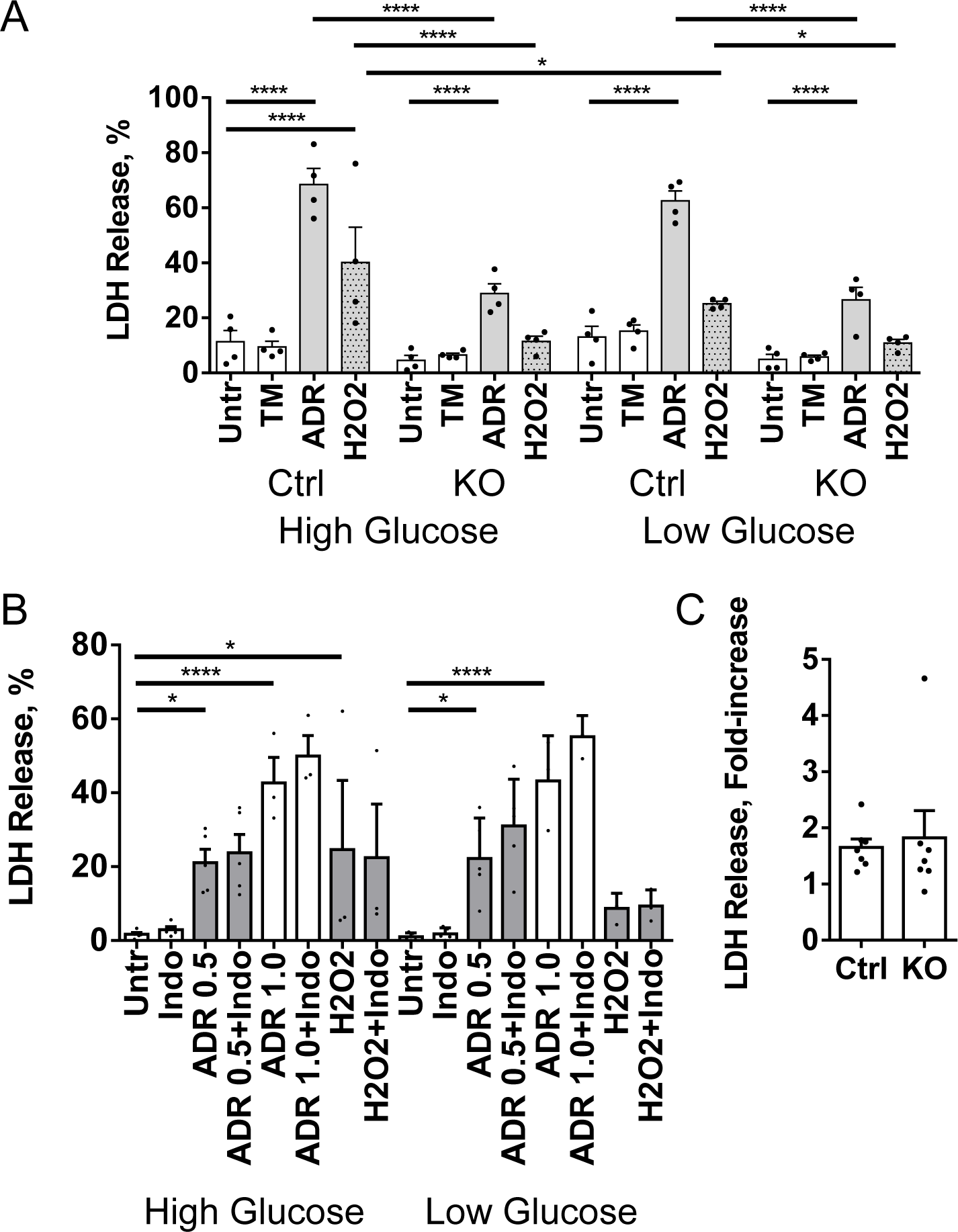
Control (Ctrl) GECs are more susceptible to injury compared to iPLA_2_γ KO GECs. A) Control and iPLA_2_γ KO GECs were untreated, or treated with tunicamycin (TM; 5 µg/ml), adriamycin (ADR; 2 µM) or H_2_O_2_ (1 mM) in high (36 mM) or low glucose (7.8 mM) media for 24 h. At the end of incubations, LDH was measured in cell supernatants and lysates, and percent LDH release was calculated. Control GECs are more susceptible to adriamycin- or H_2_O_2_-induced cytolysis compared to iPLA_2_γ KO GECs. In control GECs, H_2_O_2_-induced cytolysis is greater in high glucose. 5 experiments performed in duplicate. *P<0.05, ****P<0.0001 (ANOVA). B) Control GECs (high glucose) were untreated or treated with indomethacin (Indo; 10 µM), ADR (0.5 and 1.0 µM), H_2_O_2_ (1 mM), ADR + Indo or H_2_O_2_ + Indo for 24 h. No significant effects of Indo on LDH release are observed. 3-5 experiments performed in duplicate. *P<0.05, ****P<0.0001 (ANOVA). C) Effect of autophagy inhibition on LDH release. Control and iPLA_2_γ KO GECs were incubated with H_2_O_2_ (1 mM) in the presence or absence of 2.5 µM SBI0206965 for 24 h. SBI0206965 did independently induce LDH release (not shown). SBI0206965 augments H_2_O_2_-induced LDH release to a similar extent in control and iPLA_2_γ KO GECs. Values are fold-increase in LDH release in the presence of SBI0206965 compared to its absence (P=not significant, t-test). LDH release after H_2_O_2_ treatment in the absence of SBI020696 was 9.9±2.2% in control GECs and 4.3±0.08% in iPLA_2_γ KO GECs. 4 experiments performed in duplicate.

iPLA_2_γ was shown to induce production of prostanoids in GECs [10]. Thus, we examined if inhibition of free arachidonic acid metabolism via cyclooxygenase would affect cytolysis. Control GECs in high glucose medium were untreated or treated with indomethacin, adriamycin, H_2_O_2_, adriamycin + indomethacin or H_2_O_2_ + indomethacin. No significant effects of indomethacin on modulating cytotoxicity were observed (Fig. 6B). Glomeruli have been reported to express 12/15-lipoxygenase [37,38]. Thus, by analogy to cyclooxygenase, we tested if the 5- and 12/15-lipoxygenase inhibitor nordihydroguaiaretic acid (0.25-0.5 µM) and the 12/15-lipoxygenase inhibitor baicalein (10 µM) would modulate cytotoxicity in control GECs. Nordihydroguaiaretic acid proved to be independently toxic to GECs and could not be evaluated further. Baicalein reversed H_2_O_2_-induced LDH release in low and high glucose media, but the effect of baicalein was similar in both control and iPLA_2_γ KO GECs. Therefore, it cannot be concluded that the apparent cytoprotective effect of baicalein was associated with iPLA_2_γ.

### Production of reactive oxygen species

Reactive oxygen species may be generated in glomerular cells under diabetic conditions, and can be potentially cytotoxic [25]. We assayed production of reactive oxygen species in control and iPLA_2_γ KO GECs under hyperglycemic conditions using 2’,7’-dichlorodihydrofluorescein diacetate (DCF) fluorescence. Cells were cultured in medium containing 7.8 mM glucose. Then, medium was switched to 7.8 mM glucose + 28 mM mannitol (Man) or high glucose (36 mM) for 24 h. DCF was added and fluorescence was measured after 15 min. Although DCF fluorescence appeared to be greater in high glucose, there were no significant differences in DCF fluorescence between high and low glucose groups, nor between control and iPLA_2_γ KO GECs (Supplementary Fig. S7). The results suggest that endogenous production of reactive oxygen species, such as H_2_O_2_ is unlikely to mediate cytotoxicity in cultured GECs.

### Autophagy in cultured GECs

As shown in Fig. 5, LC3-II, reflecting autophagy, was enhanced in glomeruli of untreated and diabetic iPLA_2_γ KO mice. First, we confirmed results reported previously [7,8], showing that basal levels of LC3-II (monitored by immunoblotting) were greater in cultured iPLA_2_γ KO GECs compared to control (0.29±0.07 units in KO GECs vs 0.11±0.04 units in control; by densitometry; P=0.04, 8 experiments performed in duplicate). Next, we examined if high glucose could modulate autophagy in GECs. Cells were cultured in low glucose medium. Then, incubations were continued in low glucose + mannitol or high glucose for 24 h, and in addition, cells were treated with or without chloroquine (chloroquine blocks autophagic flux). LC3-II increased significantly after addition of chloroquine in control and iPLA_2_γ KO cells exposed to mannitol or high glucose, but there were no significant differences among these 4 groups (Supplementary Fig. S8).

Finally, we tested if inhibition of autophagy would enhance cytotoxicity. Using control GECs, we tested two compounds that have been reported to inhibit autophagy, including the ULK1 inhibitor SBI0206965 and the VPS34 kinase activity inhibitor SAR405 [39]. SBI0206965 reduced tunicamycin-induced LC3-II production, while SAR405 was not effective (Supplementary Fig. S8). Then, we incubated GECs with H_2_O_2_ in the presence or absence of SBI0206965 (we used a 2.5 µM dose, since higher doses showed some drug-dependent cytotoxicity). SBI0206965 augmented H_2_O_2_-induced LDH release in both control and iPLA_2_γ KO GECs to a similar extent (Fig. 6C), supporting the view that autophagy is a cytoprotective mechanism in GECs.

## Discussion

The present study demonstrates that deletion of iPLA_2_γ attenuates development of albuminuria in STZ-induced diabetic nephropathy. Indeed, while albuminuria in diabetic iPLA_2_γ KO mice tended to be greater than levels in the untreated iPLA_2_γ KO group, the difference was not statistically significant. iPLA_2_γ KO mice were more resistant to developing hyperglycemia compared to control mice. This result is in keeping with earlier studies, which showed that during high fat feeding, mice with global, hepatic and skeletal muscle KO of iPLA_2_γ show improved glucose tolerance and enhanced blood glucose clearance relative to controls [18,19,22]. Nevertheless, when analyzing the subset of iPLA_2_γ KO mice with greatest hyperglycemia, even this group did not show a significant increase in albuminuria, and there was only weak correlation between albuminuria and blood glucose in individual mice. Consistent with albuminuria, diabetic control mice demonstrated a higher mortality compared to the other experimental groups.

In the present study, morphologic changes in diabetic nephropathy were relatively mild. Both diabetic control and iPLA_2_γ KO mice showed increases in glomerular matrix, compared to the respective untreated groups, and the increase was greater in control mice. Diabetic control mice also showed sclerotic glomeruli and glomeruli with dilated capillary loops, in keeping with glomerular hypertension. There were, however, no significant changes in podocyte number, expression of synaptopodin, nephrin and podocalyxin, or glomerular F-actin. Decreases in synaptopodin, nephrin and glomerular F-actin have been noted previously in more robust glomerular injury models [7,32,34]. Diabetic nephropathy may be associated with inflammatory cell infiltration of the glomeruli and interstitium [25]. Interactions between resident renal cells and macrophages change the microenvironment to a proinflammatory state, contributing to tissue damage and scarring [25,35]. In our study, we did not observe significant infiltration of macrophages or neutrophils, suggesting that inflammation and inflammatory cytokines may have played only a minor role in glomerular injury.

The result that deletion of iPLA_2_γ is protective in diabetic nephropathy was not anticipated, since previously, we showed that in anti-GBM nephritis and adriamycin nephrosis, glomerular injury was actually exacerbated in iPLA_2_γ KO mice [7,8]. To examine potential mechanisms for the reduced glomerular injury in diabetic iPLA_2_γ KO mice, we studied pathways of protein degradation. In glomerulopathies, misfolded proteins may accumulate intracellularly under conditions of cellular stress. Both the ubiquitin-proteasome system and autophagy may be activated to degrade/clear these proteins [36]. There were no significant differences in protein ubiquitination among experimental groups, implying that ubiquitin-proteasome function was most likely unchanged. In contrast, autophagy (monitored by LC3-II and p62), was increased significantly in diabetic iPLA_2_γ KO mice compared to untreated and diabetic controls. Increased LC3-II in untreated iPLA_2_γ KO mice suggests that basal autophagy was enhanced in these mice. This in vivo result was recapitulated in cultured GECs, i.e. cultured iPLA_2_γ KO cells show higher LC3-II compared to control. The findings are in keeping with our earlier studies, which showed increased autophagy in glomeruli of iPLA_2_γ KO mice (both untreated and treated with adriamycin), and in resting iPLA_2_γ KO GECs in culture [7,8]. We had also shown that mitophagy is enhanced in iPLA_2_γ KO GECs [8], which may facilitate removal of the damaged mitochondria observed in these cells.

Higher autophagy in iPLA_2_γ KO mice may confer cytoprotection. This view is supported by studies in cultured GECs, where autophagy was cytoprotective. Indeed, autophagy is believed to be a major protective mechanism to clear misfolded proteins in podocytes [30,36,40]. The reason why autophagy was associated with protection from diabetic glomerular injury in iPLA_2_γ KO mice, but was insufficient to be protective in earlier studies of iPLA_2_γ in experimental glomerulonephritis may be related to the acuity of the injury. In anti-GBM nephritis and adriamycin nephrosis, glomerular injury occurred over the span of a few days to 4 weeks [7,8], while in the present study, diabetic nephropathy was a chronic lesion, developing over 5-6 months. Such distinct effects of iPLA_2_γ are perhaps not entirely surprising, since as discussed above, iPLA_2_γ activation was shown to be both protective or deleterious depending on the pathophysiological context [7,14–16,21].

There has been substantial interest in the role of autophagy in the pathogenesis of experimental diabetic nephropathy; however, conflicting results have been reported, and the role of autophagy remains to be defined precisely [29,30]. For example, in one study, the basal level of autophagy in podocytes was reduced in mice with STZ-induced diabetes [41]. In contrast, deletion of the Atg5 autophagy component in podocytes resulted in accelerated podocyte injury and albuminuria in STZ diabetic nephropathy [42]. A similar phenotype developed after deletion of Atg5 in glomerular endothelial cells. Thus, autophagy may be an important protective mechanism in both cell types of the glomerular capillary wall.

To further explore how deletion of iPLA_2_γ may have decreased albuminuria in diabetes, additional studies were conducted in cultured GECs. To recapitulate the diabetic environment, GECs from control and iPLA_2_γ KO mice were exposed to media with high glucose concentration. Control GECs were more susceptible to adriamycin- or H_2_O_2_-induced cytolysis compared to iPLA_2_γ KO GECs, and in control GECs, H_2_O_2_-induced cytolysis was greater in high glucose medium. Thus, greater cytolysis in control GECs recapitulates albuminuria and podocyte injury in vivo.

Previous studies have shown that glomerular cells contain cyclooxygenase and 12/15-lipoxygenase activities [37,38,43], and inhibition of cyclooxygenase-2 or 12/15-lipoxygenase reduced glomerular injury in diabetic models [37,38,44]. Furthermore, activation of iPLA_2_γ can lead to release of arachidonic acid and production of prostanoids in GECs [10]. Together, these results suggested that activation of iPLA_2_γ could potentially result in the generation of cytotoxic prostanoid, lipoxygenase or HETE product(s) that could injure glomerular cells. We observed that the cyclooxygenase inhibitor indomethacin did not, however, reduce cytotoxicity in control GECs. The 12/15-lipoxygenase inhibitor baicalein reversed H_2_O_2_-induced cytotoxicity, but the effect of baicalein was similar in both control and iPLA_2_γ KO GECs. Therefore, it cannot be concluded that the cytoprotective effect of baicalein was associated with iPLA_2_γ-dependent arachidonic acid release. Our results are similar to those in a recent study, which showed that hepatocytes from control mice were more susceptible to cytolytic injury compared to iPLA_2_γ KO hepatocytes. However, unlike our results, cytolysis in control hepatocytes was reduced with a lipoxygenase inhibitor, most likely due to inhibition of 12-HETE production [22]. In the future, it will be necessary to identify other factors that may be differentially regulated by iPLA_2_γ and hyperglycemia in glomeruli. For example, in mitochondria under stress, polyunsaturated fatty acyl chains in cardiolipin may first be oxidized by cytochrome c and subsequently hydrolyzed by iPLA_2_γ to serve as lipid second messengers. This may lead to the release of a variety of signaling molecules that so far are incompletely characterized [45].

There are some limitations to the present study. Deletion of iPLA_2_γ in our mice was global and not restricted to podocytes or other glomerular cells. Thus, we cannot exclude that the primary action of iPLA_2_γ in diabetic nephropathy occurred outside of the kidney. Nevertheless, development of albuminuria is typically associated with podocyte injury [3,4], and mitochondrial ultrastructural damage and autophagy in iPLA_2_γ KO mice were present only in podocytes, and no other glomerular cells or proximal tubular cells [7]. Results of studies of diabetes in cultured GECs have not been consistent. In one study, high glucose concentrations promoted autophagy [42], but in another, high glucose reduced the levels of autophagy markers [41]. Studies in cultured GECs may not accurately reflect the diabetic environment in vivo, and more complex cell culture models, e.g. co-culture of cell lines, may be required to address mechanistic questions [46]. Evaluation of iPLA_2_γ metabolic effects and autophagy in human diabetic kidneys is difficult, although autophagy genes and gene ontology pathways are upregulated in glomeruli in human diabetic nephropathy [47].

In conclusion, this and other studies support a protective role for autophagy in diabetic nephropathy, and suggest that autophagy could be an attractive therapeutic target in human disease. The precise molecular pathway(s) of iPLA_2_γ that mediate glomerular damage require further identification in future studies.

## Data availability statement

Data supporting the findings of this study are available within the article and the supplementary information.

## Author Contributions

AVC conceived and designed the research. JP, JG, JRN and HE performed experimental work. All authors analyzed the data and interpreted results of experiments. AVC wrote the manuscript. All authors reviewed and approved the manuscript.

