## Supplementary Figures for "Genetic Deletion of Calcium-independent Phospholipase A_2_γ Protects Mice from Diabetic Nephropathy"

Supplementary Figure S1

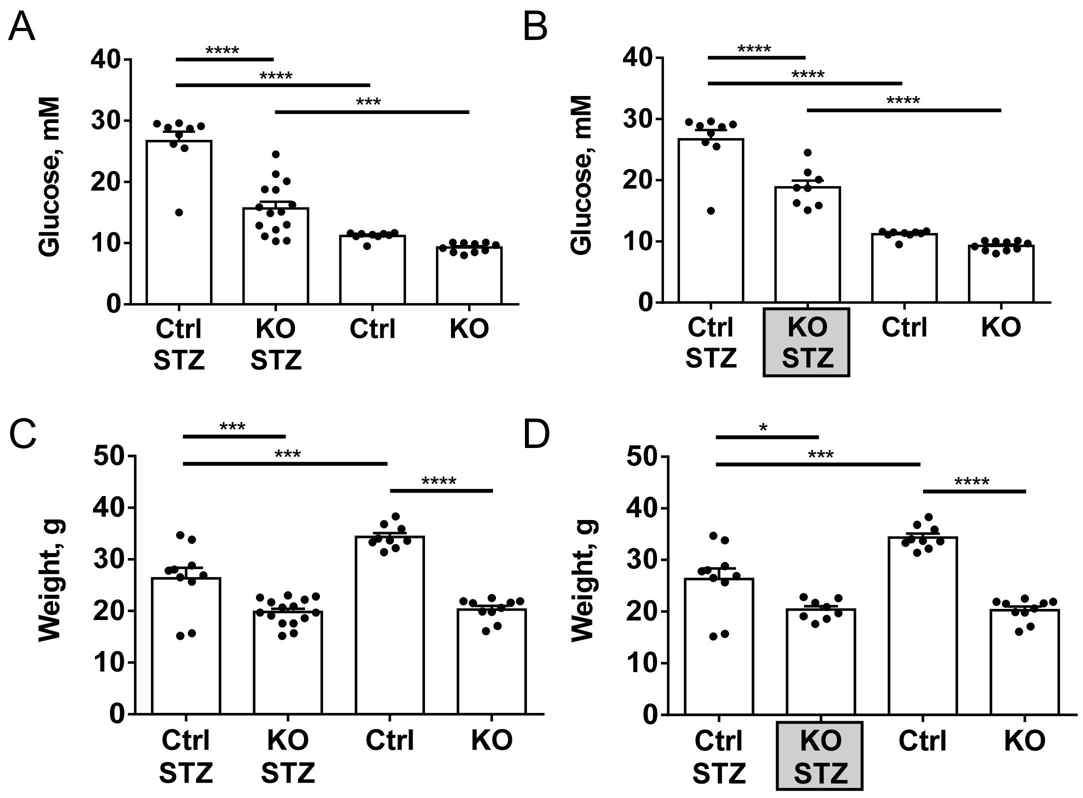

Supplementary Figure S2

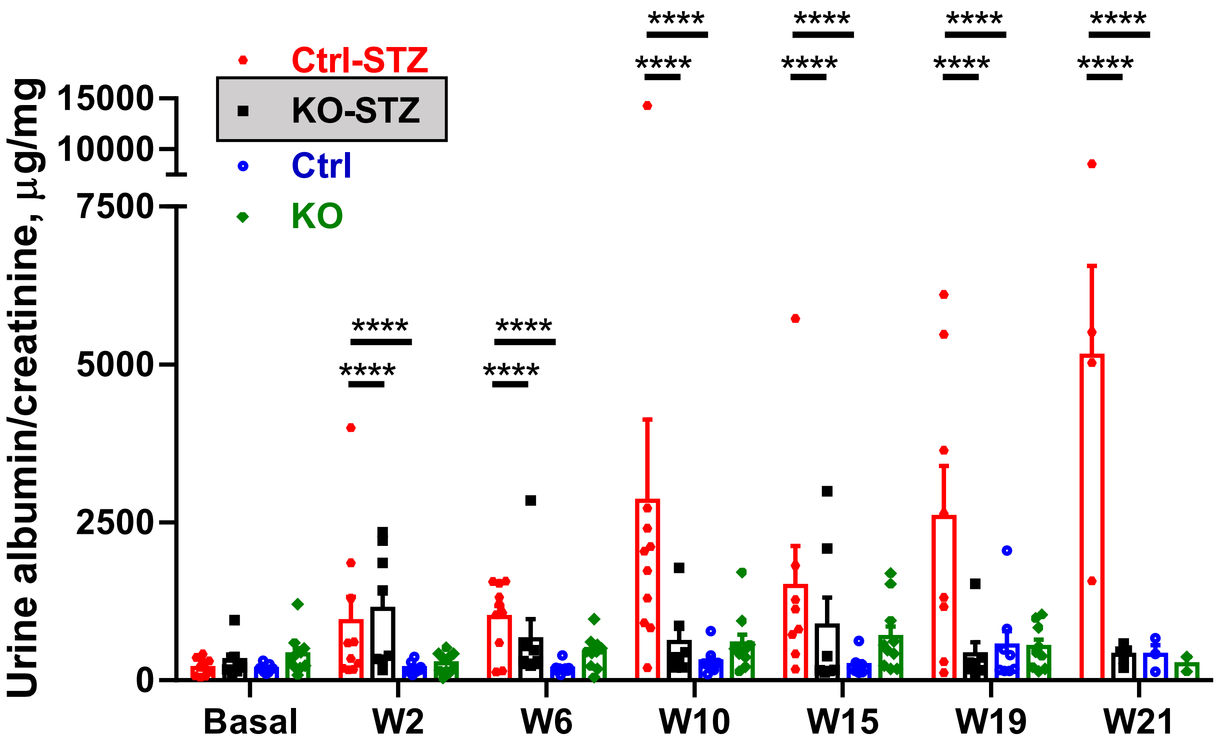

Supplementary Figure S3

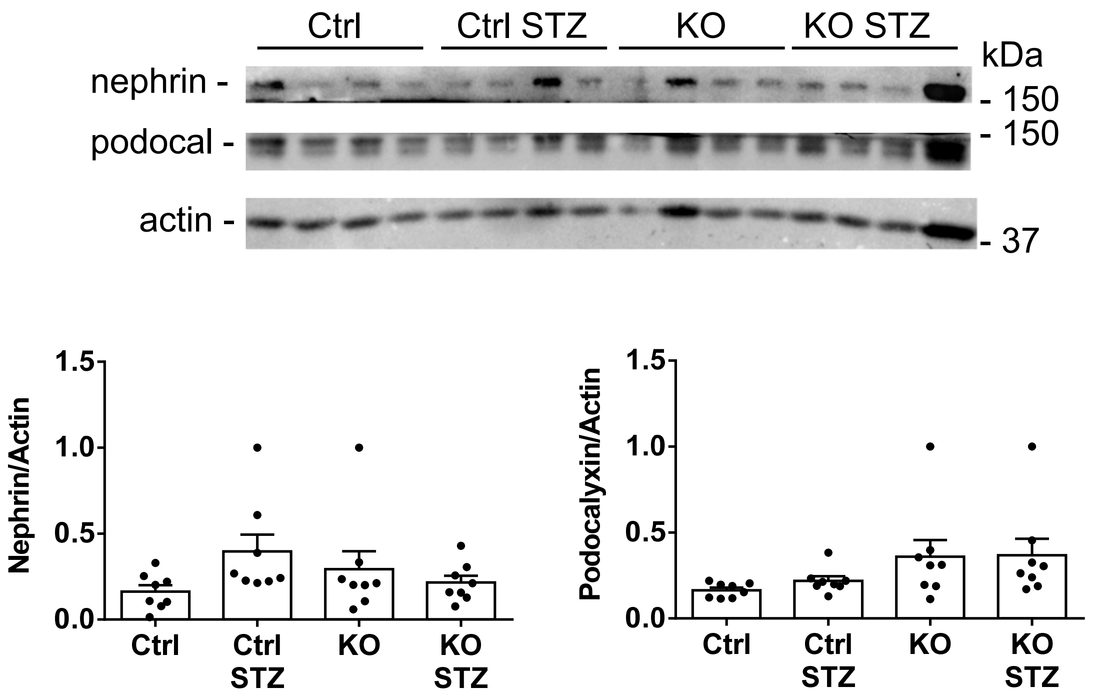

Supplementary Figure S4

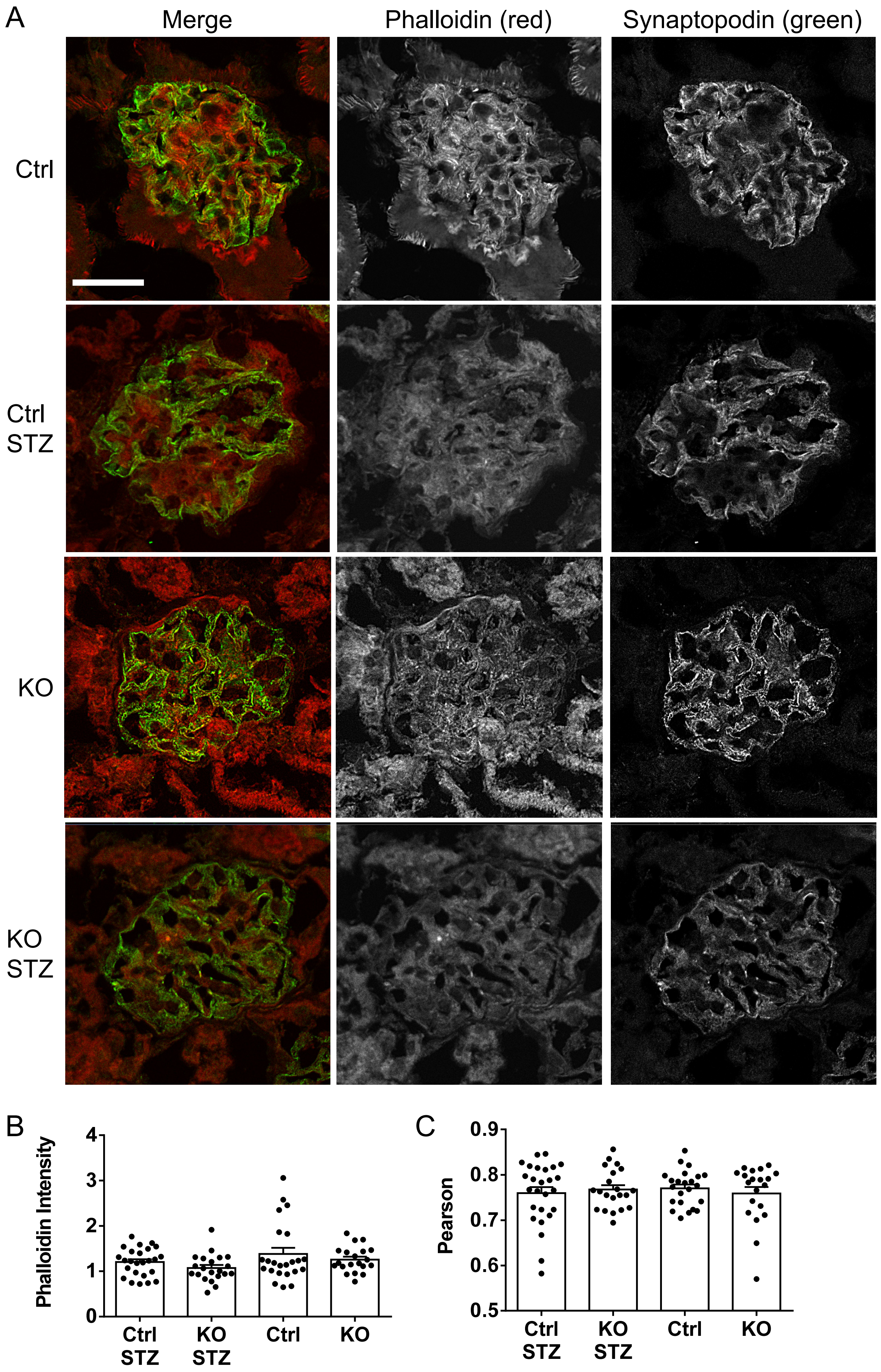

Supplementary Figure S5

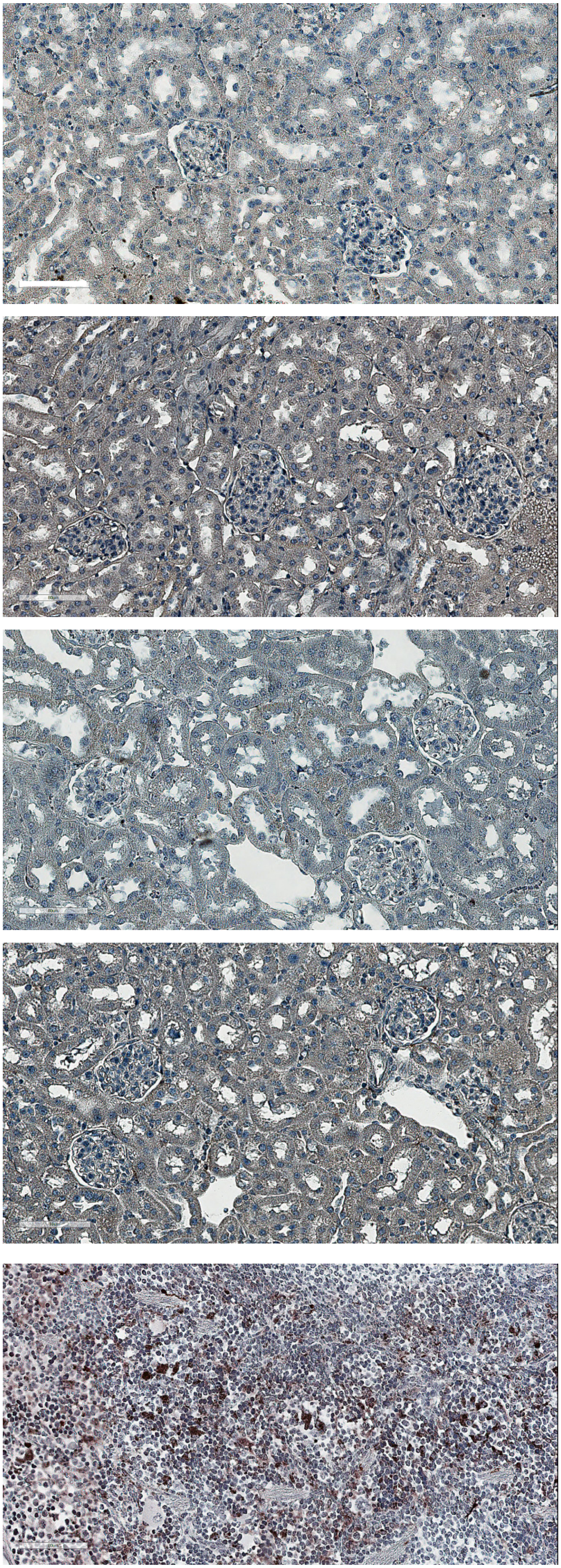

Control Untreated

KO Untreated

Control STZ

KO STZ

Spleen

Supplementary Figure S6

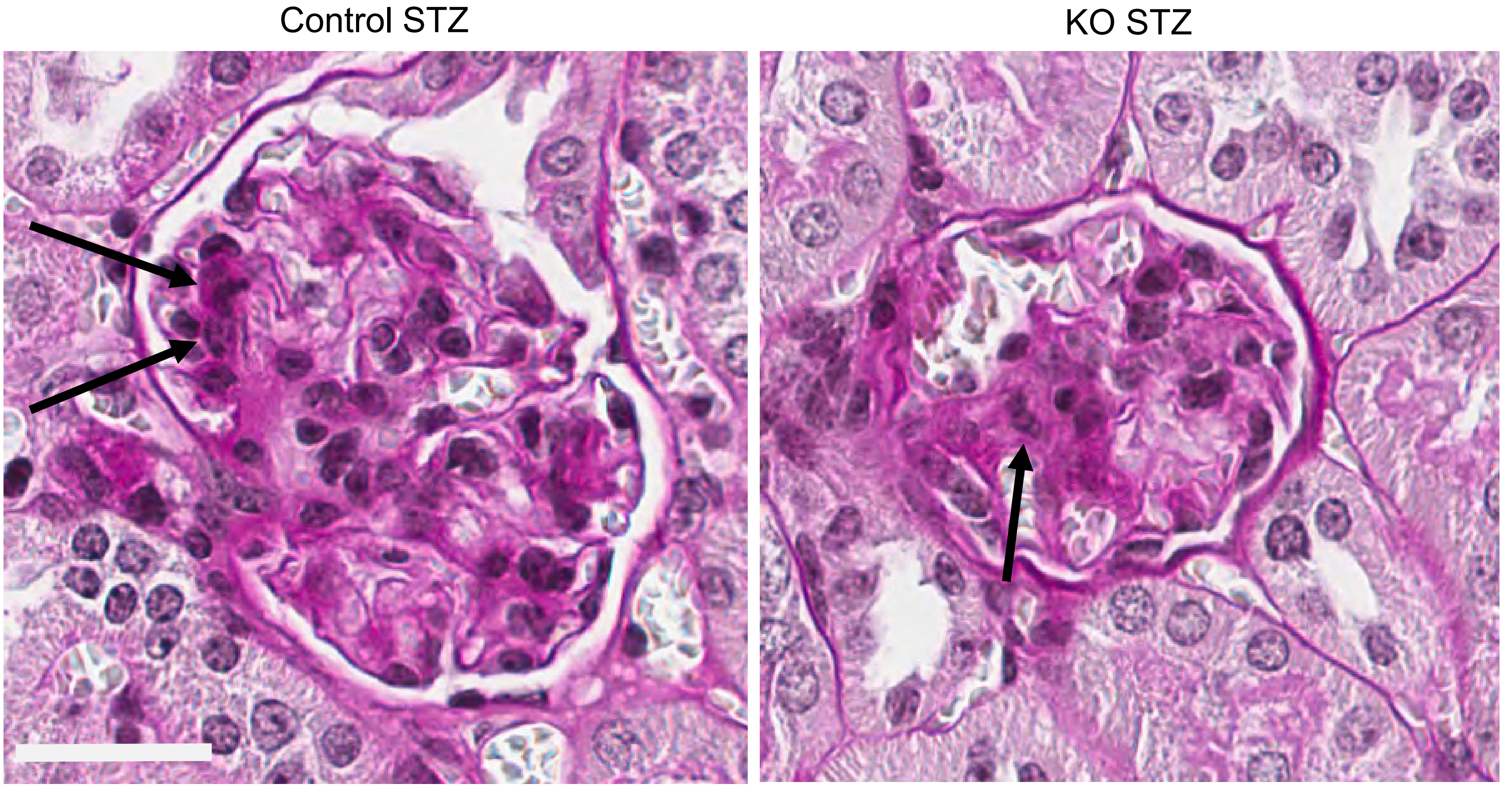

Supplementary Figure S7

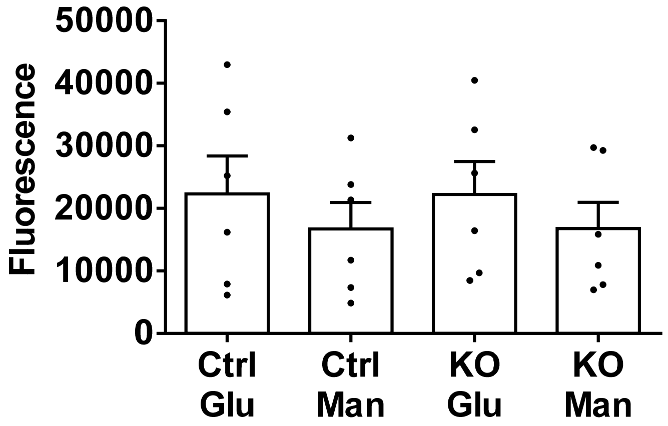

Supplementary Figure S8

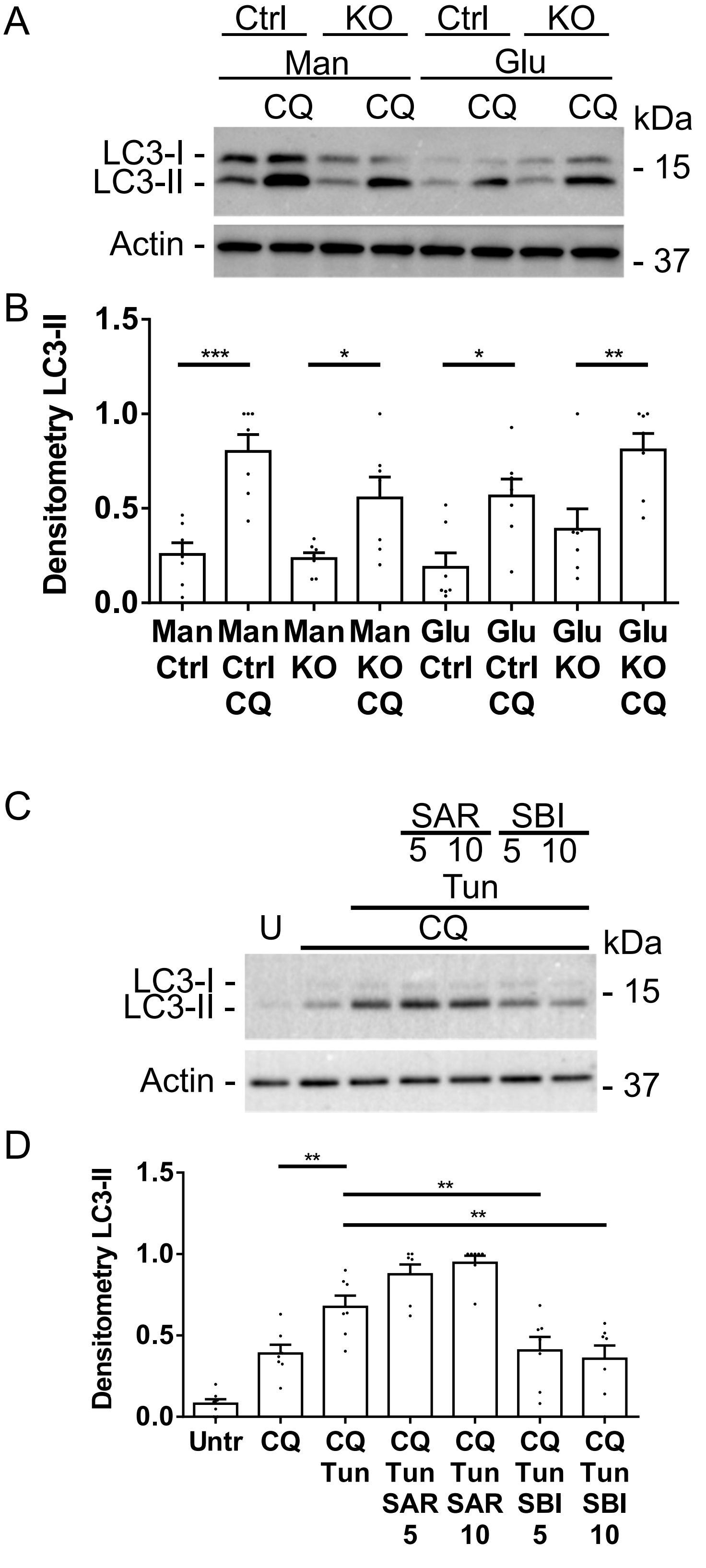
